# OM2Seq: Learning retrieval embeddings for optical genome mapping

**DOI:** 10.1101/2023.11.20.567868

**Authors:** Yevgeni Nogin, Danielle Sapir, Tahir Detinis Zur, Nir Weinberger, Yonatan Belinkov, Yuval Ebenstein, Yoav Shechtman

## Abstract

**Motivation:** Genomics-based diagnostic methods that are quick, precise, and economical are essential for the advancement of precision medicine, with applications spanning the diagnosis of infectious diseases, cancer, and rare diseases. One technology that holds potential in this field is optical genome mapping (OGM), which is capable of detecting structural variations, epigenomic profiling, and microbial species identification. It is based on imaging of linearized DNA molecules that are stained with fluorescent labels, that are then aligned to a reference genome. However, the computational methods currently available for OGM fall short in terms of accuracy and computational speed.

**Results:** This work introduces OM2Seq, a new approach for the rapid and accurate mapping of DNA fragment images to a reference genome. Based on a Transformer-encoder architecture, OM2Seq is trained on acquired OGM data to efficiently encode DNA fragment images and reference genome segments to a common embedding space, which can be indexed and efficiently queried using a vector database. We show that OM2Seq significantly outperforms the baseline methods in both computational speed (by two orders of magnitude) and accuracy.

**Availability and implementation:** https://github.com/yevgenin/om2seq

## 1. Introduction

Optical genome mapping (OGM) is a method for mapping optical images of linearly extended and labeled DNA fragments to reference genome sequences [Neely et al., 2010, Michaeli and Ebenstein, 2012, Jeffet et al., 2021]. It has shown potential in a range of applications, including the detection of structural and copy-number variations [Ebert et al., 2021], identification of DNA damage [Torchinsky et al., 2019], epigenomic profiling [Gabrieli et al., 2018, Sharim et al., 2019, Gabrieli et al., 2022, Nifker et al., 2023], and microbial species identification [Grunwald et al., 2015, Wand et al., 2019, Bouwens et al., 2020, Müller et al., 2020]. OGM’s strengths lie in its ability to image genome fragments up to megabase size and detect both large-scale structural and copy number variations, as well as working with low quantities of target DNA, which is particularly useful in cultivation-free pathogen identification [Müller et al., 2020, Nyblom et al., 2023]. Typically, OGM involves labeling DNA with fluorescent markers that bind to specific short genome sequence motifs, linearly extending the labeled DNA fragments, and optically imaging the labeled DNA fragments, followed by image analysis [Neely et al., 2010, Deen et al., 2015, Wu et al., 2018, Jeffet et al., 2021]. The mapping process to reference genome sequences uses alignment algorithms [Valouev et al., 2006, Mendelowitz and Pop, 2014, Bouwens et al., 2020, Dehkordi et al., 2021].

When it comes to mapping a labeled DNA molecule image to a reference genome, computational approaches vary based on the density of labeling. If labels are sparse enough for individual fluorescent tags to be distinguished, standard localization techniques such as emitter centroid fitting are applied to identify the labels’ positions, followed by the use of Dynamic Programming (DP) algorithms to align these positions with the expected locations of the labeled motif in the reference genome sequence [Valouev et al., 2006, Lelek et al., 2021]. Conversely, densely labeled motifs necessitate aligning the intensity profile of the imaged molecule with the theoretical intensity profile derived from the reference genome by using cross-correlation [Grunwald et al., 2015, Wand et al., 2019, Müller et al., 2020, Bouwens et al., 2020]. The DeepOM work [Nogin et al., 2023b] introduced a deep learning method for OGM, combining Convolutional Neural Network (CNN) based label localization in DNA fragment images with DP for aligning localized label positions to the reference genome.

OGM accuracy is crucial, especially for applications where the sample’s target DNA is scarce or where comprehensive genome coverage per mapping experiment is essential, such as in cultivation-free pathogen identification or detection of rare variants and epigenomic mapping [Gabrieli et al., 2018, Müller et al., 2020, Margalit et al., 2022, Gabrieli et al., 2022]. Mapping computation speed, or the mapping speed, carries equal importance, particularly when dealing with extensive DNA image datasets or when mapping to a diverse array of organism genomes for pathogen identification. Current methods are relatively slow, since for each query image, they scan the entire genome for the best match.

With those challenges in mind, we develop a novel computational method for OGM named OM2Seq, which is inspired by deep learning retrieval approaches, like Dense Passage Retrieval (DPR) [Karpukhin et al., 2020], which were initially proposed for retrieving text passages from extensive text datasets. OM2Seq is based on Transformer-based encoders [Vaswani et al., 2017] that encode both DNA molecule images and reference genome sequence segments into a unified embedding space. This enables efficient and accurate retrieval of the nearest candidate matches from a pre-computed database of genome sequence embeddings using the image embeddings.

## 2. Materials and Methods

We introduce OM2Seq, a new method for mapping DNA molecules from OGM images to the reference genome. The primary objectives of OM2Seq are to enhance the mapping accuracy, particularly for short molecules where accuracy can be challenging, and to increase the mapping speed, which refers to the speed of the computational process needed for mapping a set of DNA molecules.

The training methodology, described in detail in Section 2.5, consists of training of two encoders (Section 2.4) that encode images of DNA molecules and segments of reference genome sequences into a shared embedding space, in which distances represent the degree to which a DNA image matches a genome segment. Unlike DeepOM, which relied on simulated images for training [Nogin et al., 2023b] (and used real data only for evaluation), our training data set consists of actual microscopy images of human DNA fragments.

Following the encoder training phase, the pre-calculated embeddings of reference genome sequence segments are indexed in a vector database. During the inference phase, described in detail in Section 2.6, given an image of a DNA molecule, the Image Encoder produces an embedding for that image. This image embedding is then utilized to conduct a search within the vector database for the nearest *K* candidate reference sequence segments, with respect to some distance metric. The image query can optionally be further aligned to these retrieved candidates using a specialized OGM alignment method, such as DeepOM.

### 2.1. Data acquisition

For the purpose of training, validation, and testing in this work, we utilized a dataset composed of images featuring linearly extended human DNA fragments. These images were captured with the use of the Bionano Genomics Saphyr instrument and Saphyr chips (G1.2). A total of 100,000 DNA molecule images were used, all of which have been made publicly available as detailed in Section 4. An illustrative example can be found in Figure 1. Individual molecules were cropped from captured raw images using the segmentation coordinates in the Bionano software’s output BNX files. The comprehensive protocol for the DNA extraction and labeling process is described in previous work [Nogin et al., 2023b].

**Fig. 1:**
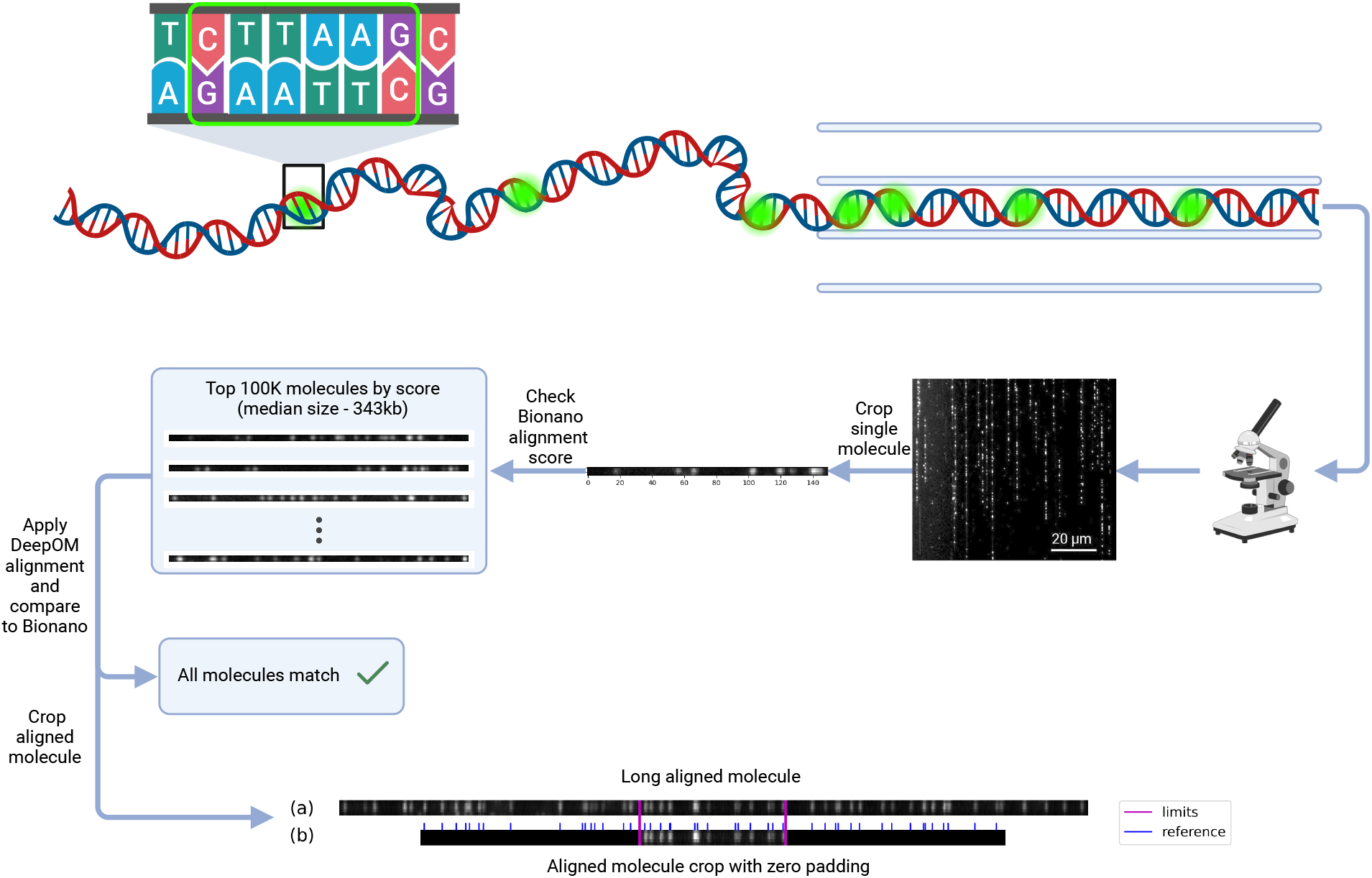
Training and validation data. The training data, as detailed in Section 2.3, was generated from images of DNA molecules acquired with the Bionano Saphyr instrument, labeled at a specific sequence pattern (CTTAAG, see Section 2.1). Individual molecule images were extracted using the output of the Bionano image processing pipeline. For the creation of the ground-truth set, a set of molecules with high Bionano alignment confidence score were chosen and their alignment was validated with the DeepOM algorithm [Nogin et al., 2023b]. From this ground-truth set, subsets for training, validation, and test were sampled. Shown are: (a) an example of a long DNA molecule image in the ground-truth set and (b) an example cropped fragment zero padded to a specific length (used later for training or evaluation), with its crop limits shown, alongside the corresponding reference genome segment with the labeled pattern positions shown.

In brief, DNA was extracted from approximately one million cells from the U2OS human cell line, following the Bionano Prep Cell Culture DNA Isolation Protocol developed by Bionano Genomics. For the labeling step, one microgram of DNA was processed using the Bionano Genomics DLS labeling kit in conjunction with the DLE-1 enzyme, which labels the genome sequence CTTAAG. The reaction mixture, with a total volume of 30 microliters, contained 6 microliters of 5x DLE-buffer, 2 microliters of 20x DL-Green, and 2 microliters of the DLE-1 enzyme, all supplied by Bionano Genomics. This mixture was subjected to an incubation period of 2 hours at a temperature of 37 degrees Celsius.

### 2.2. Genome sequence data

Human genome GRCh38.p14 from https://www.ncbi.nlm.nih.gov/datasets/ was used.

### 2.3. Training dataset

The training dataset for OM2Seq was compiled from the collection of acquired images of DNA molecules on the Bionano Genomics Saphyr platform (Section 2.1), and a selection for the ground-truth set of the top 100,000 molecules out of roughly 25.4 million was made based on the alignment confidence scores provided by the Bionano software in the XMAP output files. Typically, the acquired images are 5 pixels wide and extend to roughly 1000 pixels in length. The lengths of the selected molecules ranged from a minimum of 181 kilobases (kb) to a maximum of 768 kb, with a median size of 343 kb. Selecting such long molecules ensures a strong ground-truth for the training data, as the probability of alignment error for these long molecules is negligble, as was shown empirically and theoretically in terms of OGM Information Theory [Nogin et al., 2023a]. The distribution of molecule lengths and alignment confidence scores in the ground-truth set, and the genome coverage of those molecules are shown in Supplementary Section S2.

Each DNA molecule image was aligned with the human genome (GRCh38.p14 release) by employing the DeepOM method [Nogin et al., 2023b]. Given that DeepOM achieves nearly 100% mapping accuracy for the long molecules selected for this dataset, these alignments serve as a reliable ground-truth. Additionally, these DeepOM alignments were compared and validated to be the same as results from Bionano’s aligner.

The dataset was divided into subsets designated for training, validation, and testing, with both the validation and test sets comprising 1,000 molecules each.

To prepare OM2Seq for effective mapping accuracy, even on molecules as short as 30kb, the data was manipulated by cropping shorter fragments from the longer original DNA molecule images (see Figure 1). For training, a batch of *n* cropped fragments is generated at each step, accompanied by their respective reference genome segments (see Figure 3).

A training batch is assembled in the following manner: *n* aligned molecules are randomly chosen from the training set. For each of these molecules, a random cropping length, sampled with log-uniform distribution in the range 30kb to 200kb, and a random starting position are selected. This process yields a cropped fragment image and the corresponding ground-truth reference position within the original molecule (whose alignment to the reference genome is known), as well as the matching genome sequence reference segment. The validation batch creation follows this process using molecules from the validation set.

### 2.4. Model Architecture

Our model architecture takes cue from the Transformer encoder utilized in the WavLM work [Chen et al., 2022], which was initially developed for learning speech representations from extensive unlabeled speech data and achieved state-of-the-art results on various speech-related downstream tasks. The decision to leverage this approach is anchored in the analogy between speech, represented as an analog, noisy, time-series audio signal encoding textual information, and OGM images of DNA molecules, which can be seen as one-dimensional intensity signals encoding genome sequences.

Incorporated within our architecture is a convolutional feature encoder followed by a transformer encoder, as depicted in Figure 2. The first transformer output (CLS token as in BERT [Devlin et al., 2019] and in DPR [Karpukhin et al., 2020]) in the output sequence is taken as the output embedding vector, while the other outputs are ignored (a common practice in those kinds of models, as done in BERT and DPR, and a similar result could be achieved by averaging the outputs). The OM2Seq model is composed of two Transformer-encoders: one dubbed the Image Encoder, tasked with encoding DNA molecule images into embedding vectors, and another called the Genome Encoder, devoted to transforming genome sequence segments into their embedding vector counterparts. Starting from the base WavLM architecture, for the Image Encoder, in the first convolutional layer the number of input channels was set to the image width in pixels (5), the kernel size to 4, and the stride to 1. For the Genome Encoder, the input genome reference label positions vector is binned into 320bp bins, and the number of positions in each bin is counted. This counts vector is the input to the first convolutional layer, which has only one channel, and the same kernel and stride as the Image Encoder. For the transformer, 3 hidden layers and 6 attention heads were used for both encoders. The transformer hidden size, intermediate size and the convolutional hidden dimension were all scaled down by a factor of 0.072 with the other parameters used as WavLM’s defaults. The chosen model down-scaling factor was the one giving best validation accuracy, after multiple training runs. In result, each embedding vector produced by the encoders has an embedding dimension of 96 and the model has 651,461 parameters.

**Fig. 2:**
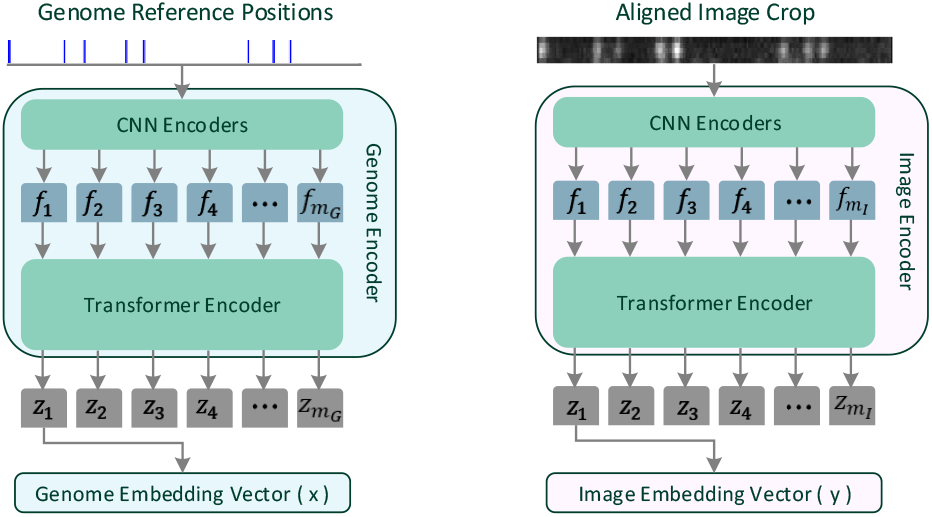
Model architecture. The model of OM2Seq, as detailed in Section 2.4, is built upon a Transformer encoder architecture, which processes images of DNA molecules and reference genome sequence segments into a unified embedding space. This design is based on the architecture of WavLM [Chen et al., 2022], featuring a convolutional feature encoder (with outputs *f*_*i*_) followed by a transformer encoder (with outputs *z*_*i*_). The number *m* (*m*_*G*_ for the Genome Encoder, *m*_*I*_ for the Image Encoder), of extracted feature vectors *f*_*i*_, is dependent on the input length, and the CNN stride parameters. The first transformer output, *z*_1_, in the output sequence is taken as the output embedding vector, and the others are ignored.

### 2.5. Model Training

For the training of the embeddings, we implemented the contrastive loss function used in CLIP [Radford et al., 2021]. CLIP’s training regime utilizes a vast dataset of image-text pairs, to encode images and their captions to a common embedding space. For each training batch of size *n*, a cosine similarity matrix *s* is constructed from the dot products of image and text embeddings. Consequently, this produces a *n × n* matrix for the batch, which forms the basis for calculating the contrastive loss.

Cosine similarity is calculated as the dot product of two vectors, normalized by the product of their magnitudes. In an ideal scenario where embeddings are perfect, the similarity matrix would be the identity matrix. Therefore, the identity matrix serves as the target for optimizing the similarity matrix. This is achieved by applying a symmetric cross-entropy loss function to both rows and columns of the similarity matrix with the identity matrix as target, as shown in Figure 3. The similarity matrix is also scaled by a learnt factor *τ* initialized to 0.07 as in the CLIP work, and the loss function is expressed as:

**Fig. 3:**
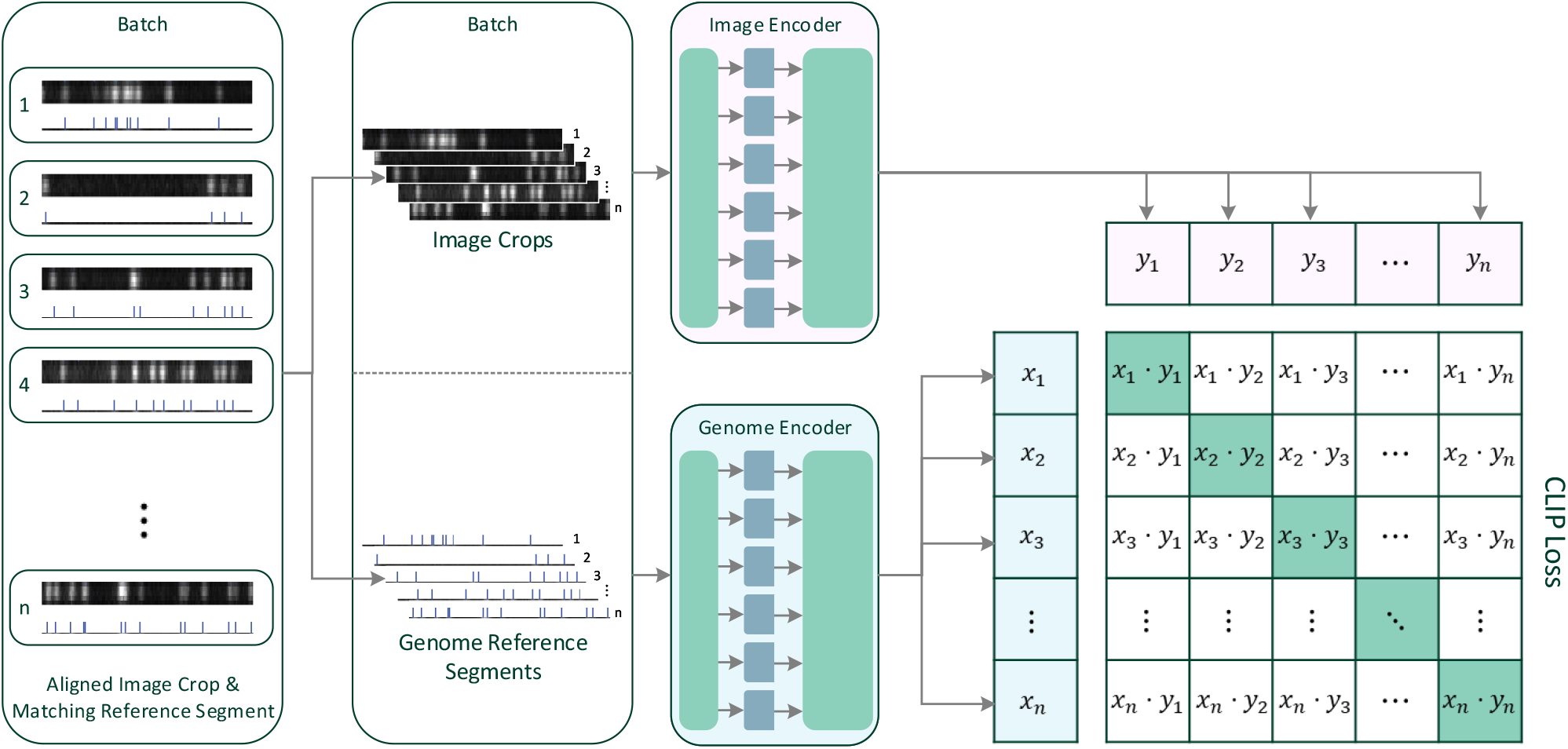
Training the model. As detailed in Section 2.5, the model training process involves encoding images of DNA molecules and reference genome sequence segments into a unified embedding space and trained using a contrastive loss function, similar to CLIP [Radford et al., 2021]. Both the Image Encoder and the Genome Encoder architectures are detailed in Figure 2. All images are zero-padded to the same constant size during training and inference. The reference genome segments are always taken with a constant length.

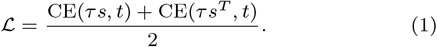

where *s* represents the similarity matrix (and *s*^*T*^ its transpose, to enforce the identity target both on rows and columns), *t* is the target identity matrix, and CE denotes the cross-entropy loss function, defined as:

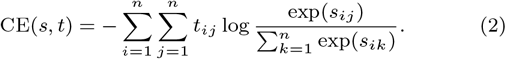

Since *t* is the identity matrix, the cross-entropy function can be simplified to:

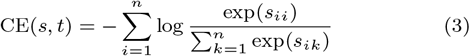

In the context of our OGM data, each batch comprises *n* pairs of cropped DNA molecule images and their respective genome reference segments, as outlined in Section 2.3. By denoting the normalized genome embedding vectors as *x*_*i*_ and the normalized image embedding vectors as *y*_*j*_, we calculate the element of the cosine similarity matrix as *s*_*ij*_ = *x*_*i*_ *· y*_*j*_.

The minimization of the loss function was done via stochastic gradient descent using the AdamW optimizer [Loshchilov and Hutter, 2019], across 131,600 steps, with learning rate 2 *×* 10^*−*4^, batch size 256, weight decay 1 *×* 10^*−*2^, and betas (0.9, 0.999). Additionally, validation loss was assessed using a validation batch at intervals during the training process (see Supplementary Figure S1). The training was conducted on a single NVIDIA A100 GPU, and took around 40 hours.

### 2.6. Inference and retrieval

For the retrieval of genome sequences based on OGM image queries, we adopted techniques from the Dense Passage Retrieval (DPR) work [Karpukhin et al., 2020], which originally trained models to encode text passages and questions into embeddings used for efficient retrieval of passages for open question answering. In DPR, a contrastive loss function similar to CLIP [Radford et al., 2021] was used to train embedding encoders. These encoders were trained on a dataset of passages from Wikipedia articles, and the embeddings were stored in a FAISS vector database [Johnson et al., 2019].

In our application for OGM, the same approach was used, with reference genome sequence segments as reference passages and DNA molecule image embeddings serving as queries. For efficient retrieval, we pre-computed the reference genome sequence embeddings and stored them in a FAISS vector database, as in DPR (Figure 4). The reference embedding database was created by taking sequential segments of the human genome with a length of 200 kb and offsets starting in increments of 30kb, resulting in approximately 10^5^ segments. The trained Genome Encoder model (Section 2.5) then generated embeddings for each segment, which were stored in the FAISS vector database.

**Fig. 4:**
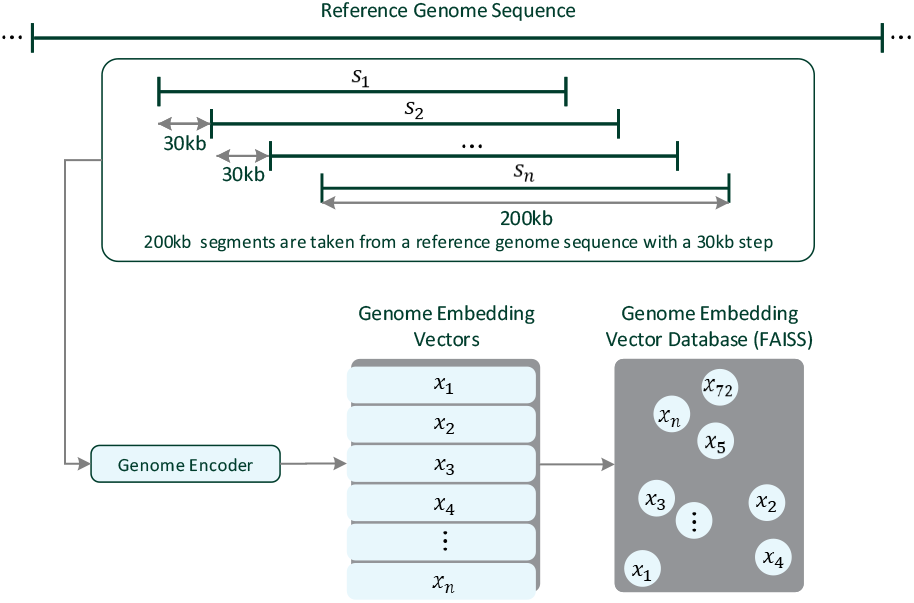
Pre-computed Genome Vector Database. As detailed in Section 2.6, the genome vector database, later queried in the inference phase, is pre-calculated by applying the trained Genome Encoder on 200kb genome segments (*s*_*i*_) extracted from a long genome reference with 30kb offsets. The embedding vectors (*x*_*i*_) are then indexed into a FAISS vector database for fast retrieval [Johnson et al., 2019].

At inference time (Figure 5), an image of a DNA molecule is processed by the Image Encoder model to produce an image embedding. This embedding is queried against the vector database to retrieve the nearest (by cosine-similarity) *K* candidate genome reference segments. The database returns these *K* candidates, complete with their embeddings and reference genome offsets. Candidates are ranked according to similarity to the query, with the highest similarity indicating the predicted match to the reference sequence. This retrieval process is much faster than alignment based methods since the indexing structure of the vector database allows for reduced computational complexity, which can be logarithmic as a function of the reference genome length (and the number of embedded segments) [Johnson et al., 2019], as opposed to linear in the case of alignment based methods.

**Fig. 5:**
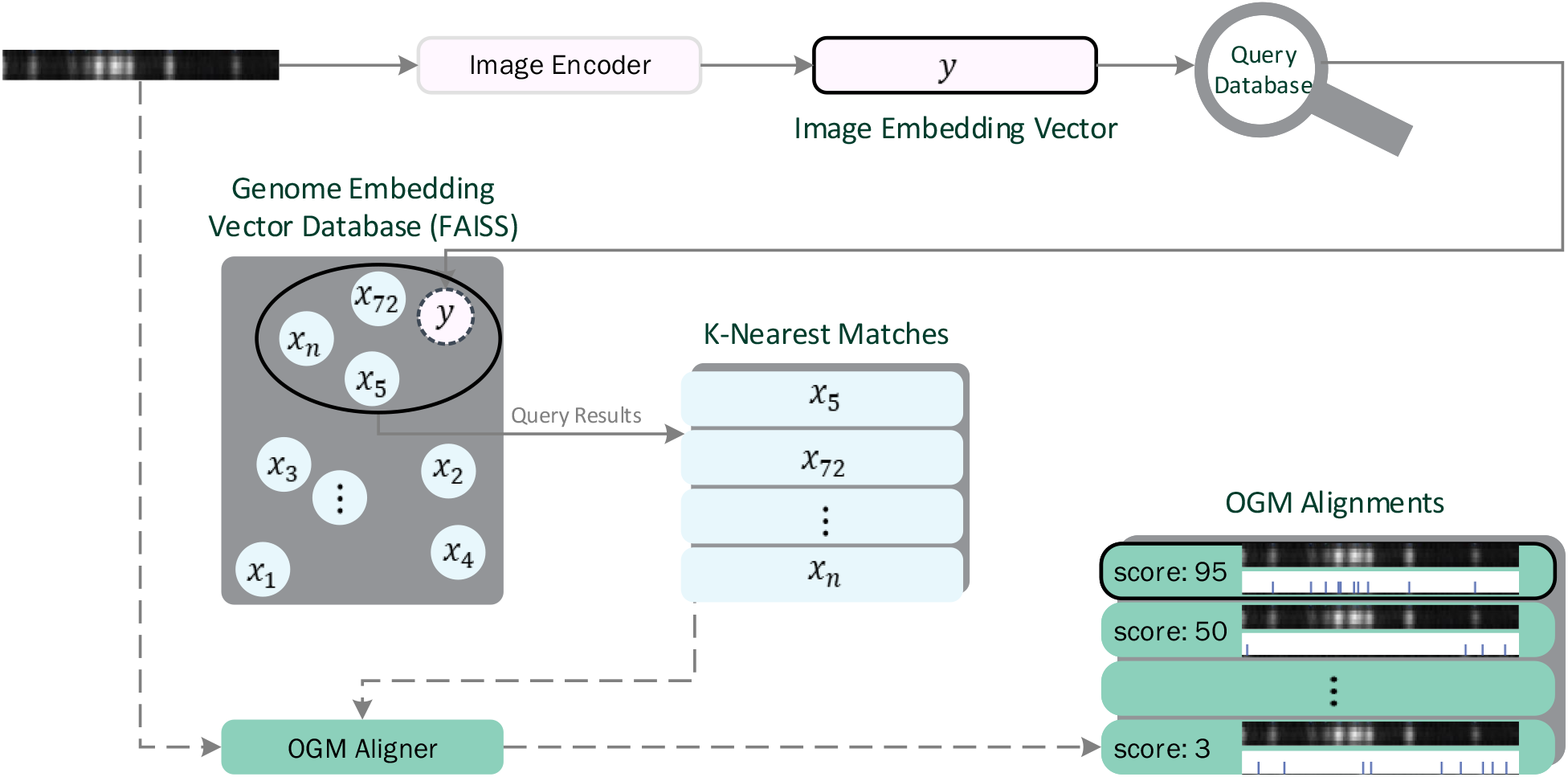
Inference and retrieval. As detailed in Section 2.6, the inference and retrieval process, inspired by DPR [Karpukhin et al., 2020], involves encoding an image of a DNA molecule to an embedding vector (*y*) and retrieving the nearest *K* candidate reference sequence segments from a pre-computed vector database of their embeddings (*x*_*i*_, see Figure 4). In a final optional step, an OGM aligner can be used to precisely align the nearest *K* matched reference segments (of size 200kb) with the molecule image (which could be as short as 30kb), and choose the highest scoring alignment. This way both high accuracy and high computation speed can be achieved (Section 3).

To enhance mapping accuracy, the nearest *K* candidate reference sequence segments can be aligned to the query image using an OGM alignment algorithm, such as DeepOM [Nogin et al., 2023b]. The best performing candidate in terms of alignment score is then selected as the predicted mapping. This process leads to both a speed increase in the mapping speed and an improvement in mapping accuracy (Figure 5).

### 2.7. Baseline and Evaluation

We use DeepOM, which also contributed to the training data generation as described in Section 2.3, as a baseline for comparison. DeepOM is a two-step method, first localizing labels on the DNA fragment image with a CNN, and then aligning the fragment to the genome with a DP algorithm.

Our evaluation comprises assessing the accuracy of OM2Seq in comparison with DeepOM, as well as measuring the mapping speed (computational speed) of both methods. Additionally, we examine an integrated approach where OM2Seq retrieves the nearest *K* = 16 candidate reference sequence segments, and DeepOM aligns the query image to these candidates, utilizing its alignment score to select the best match.

For each tested DNA fragment length, with length ranging from 30kb to 200kb (their lengths in basepairs are approximated from the alignment to the genome reference of the molecule from which they are cropped), we generate *N* = 1000 queries from cropped DNA molecule fragments, each derived from a distinct molecule in the test set, as aforementioned in Section 2.3. Each query has a known ground-truth reference position. The selected OGM mapping method maps each query, and the resulting mapping is compared to the ground-truth reference coordinates.

An overlap between predicted and ground-truth reference segments deems a prediction correct. Since we have a ground-truth alignment for each query image, we know the genome reference start and stop positions for this query. A segment overlap (intersection) is computed between the query genome segment and the predicted genome segment, and a non-zero overlap is considered correct. Accuracy is then defined as the ratio of correct predictions to the total number of queries. It is important to note that given a short image query of say 30 kb, the method provides its position up to a resolution of the predicted 200 kb genome reference segment. To obtain a more exact position, an additional alignment method should be applied, as described earlier.

In addition to the accuracy, we recorded the time taken to map the entire set of *N* queries by each OGM method and calculated the mapping speed as the sum of the lengths of the queries in base pairs divided by the runtime. The evaluation was performed on a Google Cloud instance with 12 vCPU cores and an NVIDIA A100 GPU.

## 3. Results

Our experimental process included generating a dataset of aligned DNA molecules as delineated in Section 2.3, training the OM2Seq model per the methods outlined in Section 2.5, with the training progress exhibited in Figure S1, and evaluating the accuracy and mapping speed of OM2Seq, DeepOM, and their combined applications on the test-set (molecules not used for training) as specified in Section 2.7. The combined application of OM2Seq and DeepOM is detailed in Section 2.6, where *K* = 16 candidate reference segments are retrieved by OM2Seq and aligned by DeepOM. For OM2Seq alone, the number of retrieved candidates was set to *K* = 1. The effect of the choice of *K* is elaborated in Supplementary Section S4. The findings regarding accuracy and mapping speed are visually presented in Figures 6 and 7, respectively. It can be seen in Figure 6 that OM2Seq in itself has comparable accuracy to DeepOM, while there is a slight accuracy drop for OM2Seq at 30kb and 200kb, which is due to the fact the training was done with crop lengths sampled from 30kb to 200kb in length, which reduced the amount of training data for these edge lengths (since the sampling distribution was log-uniform, see Section 2.3). Both methods significantly outperform the benchmarked commercial OGM alignment software provided by Bionano Genomics Inc. (which was evaluated on the same OGM dataset as here). When combining OM2Seq and DeepOM (as described in Section 2.6), the accuracy is significantly improved, for example for 50kb fragments, from 63% for DeepOM to 78% for the OM2Seq+DeepOM combination (while less than 30% for the commercial OGM aligner).

**Fig. 6:**
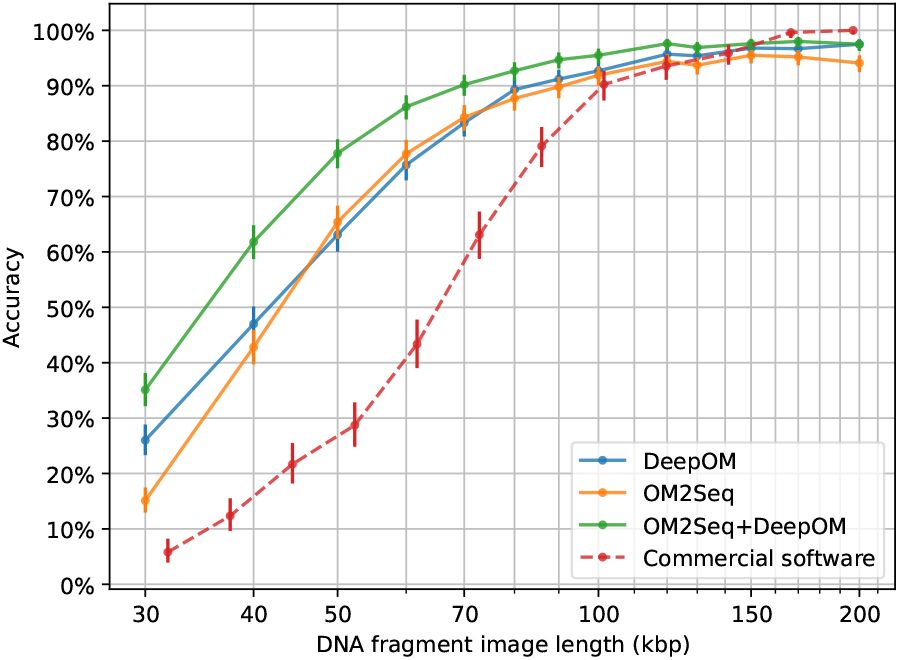
Accuracy. The accuracy of OM2Seq (number of candidates *K* = 1), DeepOM, and their combination (*K* = 16) is evaluated as detailed in Section 2.7, for various DNA fragment lengths. Accuracy results for the commercial software from the benchmark done in the DeepOM work are also shown for comparison (adapted from Figure 4b, Bionano localizer + Bionano aligner, in [Nogin et al., 2023b]). Accuracy is measured as the proportion of queries where the predicted mapping overlaps with the correct genome reference positions. Error bars indicate 95% confidence intervals, calculated utilizing the Clopper-Pearson Beta Distribution method [Clopper and Pearson, 1934].

**Fig. 7:**
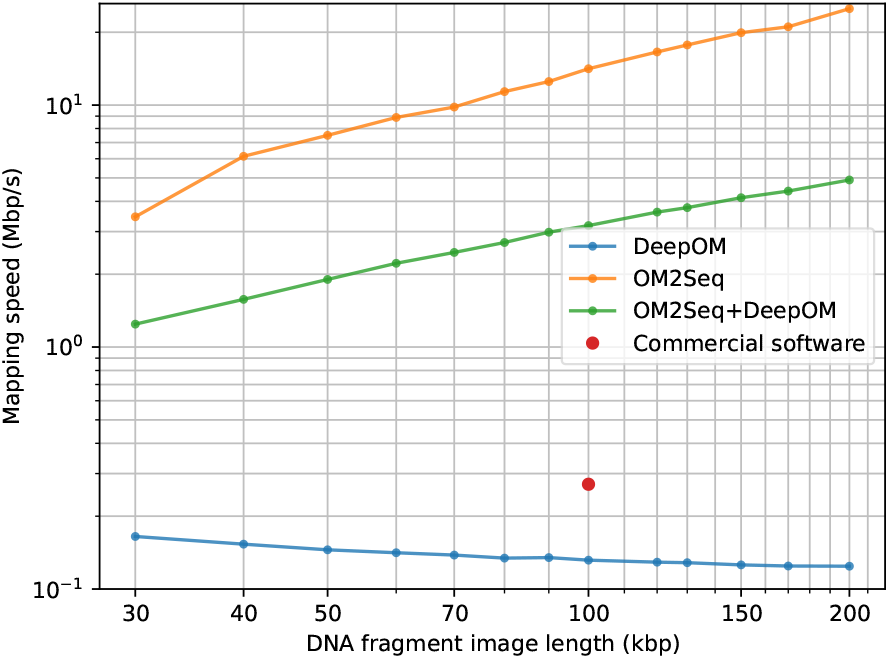
Mapping speed. The mapping speed (computation speed, as described in Section 2.7) of OM2Seq, DeepOM, and their combination for various DNA fragment lengths. It is computed as the cumulative length of the DNA fragment queries in base pairs divided by the runtime. The mapping speed of the commercial software reported in supplementary information of DeepOM [Nogin et al., 2023b] is also shown.

Beyond its improved accuracy (when used in combination with DeepOM), the great power of OM2Seq becomes apparent when comparing computation speed-up (Figure 7). In comparison to the baseline DeepOM method, our OM2Seq model demonstrated significantly faster mapping speeds (by two orders of magnitude for 200kb fragments for OM2Seq alone), meaning it could process the same genome coverage in a shorter amount of time. It should also be noted that the combination of OM2Seq and DeepOM is faster than DeepOM alone due to the small set of candidate reference segments retrieved by OM2Seq, significantly reducing the total reference sequence size to which DeepOM needs to align the images.

## 4. Discussion

The results shown here for OM2Seq have highlighted its potential to significantly advance OGM. As an approach based on training with real acquired data (as opposed to training on simulated data in DeepOM), OM2Seq boasts the significant advantage of bypassing the need for simulation-based image generation, a process that traditionally requires substantial customization to ensure the simulator output adequately reflects real images. This methodological shift allows for direct training on available real data and affords superior accuracy for mapping shorter molecules by learning to extract as much information as possible from the images, utilizing the longest molecules within a given dataset for training.

Furthermore, OM2Seq has the potential to facilitate OGM for labeling methods whose labeling mechanisms are not fully understood and for which there is no available genome alignment method. This can be implemented when two different labeling methods are employed, each with its distinct fluorescent color channel [Jeffet et al., 2021]. Having an OGM alignment method for just one of the channels, enables ground-truth alignments for both channels (assuming they are optically aligned). In addition, those ground-truth alignments enable the generation of a training dataset for OM2Seq on the second channel, for which there is no alignment method.

However, OM2Seq has the following limitations. The model requires substantial quantities of experimental data for training, which is not always easy to obtain, and while a moderate degree of accuracy might be attainable with a smaller dataset, the accuracy presented in this study is contingent on the use of data from an order of 10^5^ molecules. Furthermore, the training data could be biased since it is composed solely of long molecules that the DeepOM or Bionano alignment methods could align, cropped to shorter fragments. This was necessary since no other ground truth was available for training. Moreover, the ground-truth set was limited to molecules having high Bionano alignment confidence scores, but as we demonstrate in Supplementary Section S3, the accuracy doesn’t seem to be dependent too much on these scores. Due to these ground-truth limitations, it is difficult to assess the model’s performance on independently acquired short molecules without having some kind of ground truth for them. Such ground-truth could potentially be obtained, either by using short molecules from a known short reference sequence (say a virus for example), or adding another labeling channel on top of them, which could serve as a ground-truth validation. Another direction that could be explored is using OM2Seq to fine-tune itself. This could be done by first applying OM2Seq inference and alignment to medium-sized molecules, of say 100kb (which is still very accurate, as seen in the results Figure 6), then generating a new training dataset based on cropping those molecules as described in Section 2.3, and fine-tuning the OM2Seq model on that dataset.

It should also be noted that OM2Seq’s outputs are mappings to discrete genomic segments (of 200kb size in this study) that were indexed in the vector database as described in Section 2.6. For precise pixel-level alignment, supplementary methods such as DeepOM are still needed as a post-processing step. For future work, an additional transformer encoder-decoder architecture could be developed for this purpose and trained on the same dataset, harnessing the cross-attention mechanism of the transformer architecture [Vaswani et al., 2017] for direct alignment of the image pixels to genome reference positions, similar to transformer-based alignment of nucleotide or protein sequences [Dotan et al., 2023].

An interesting open question is - how low can we go? namely, how short can molecules be and still be aligned correctly. The label density in the ground-truth set (including the training, validation and test set) was estimated to be around 5481 bp / label, so, for example, a typical 30kb fragment (evaluated for in Figure 6) can have 5 to 6 labels. If one would have perfect label localization accuracy, this for sure would be enough to place the fragment [Nogin et al., 2023a]. OM2Seq is able to gain much better accuracy than previous methods because it works directly on images, skipping the localization step with its associated errors. In conclusion, OM2Seq showcases a method for mapping DNA molecules from OGM images to the reference genome by encoding images of DNA molecules and reference genome sequence segments into a common embedding space. The results highlight OM2Seq’s ability to achieve higher accuracy and faster mapping speeds compared to the baseline method, DeepOM. Increased accuracy on shorter molecules also implies higher genome coverage from a given sample, ultimately leading to higher diagnostic sensitivity. The increased mapping speed with OM2Seq allows for quicker analysis, which is valuable when dealing with high coverage OGM mapping of the human genome, or when mapping against large datasets of organism genomes, such as in the identification of pathogens.

## Data and Code Availability

The data and code used in this work is available on Zenodo: https://doi.org/10.5281/zenodo.11076514

## Acknowledgements

This work was supported by the Google Cloud Research Credits program with the award GCP19980904. Some figures in this paper were created with BioRender.com.

