## Supplemental Information for "OM2Seq: Learning retrieval embeddings for optical genome mapping"

April 27, 2024

#### S1 Training details

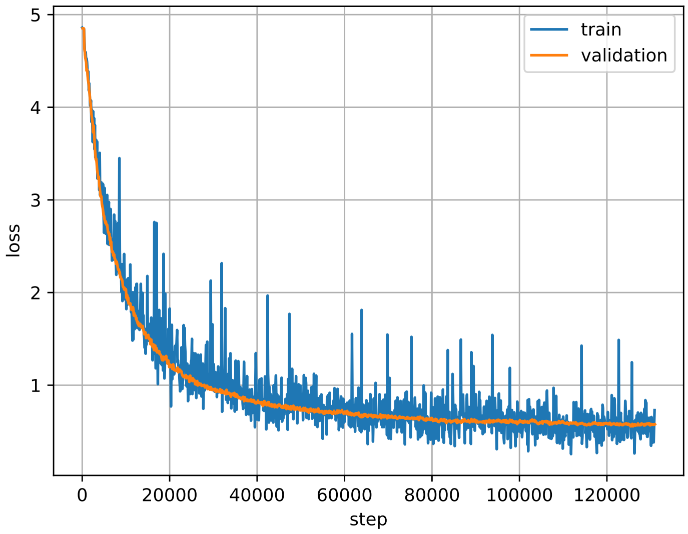


Figure S1: **Training loss.** As detailed in subsection 2.5, the training process involved minimizing the contrastive loss function, as shown in this figure. The validation loss is calculated using a validation batch at intervals during the training process.

### S2 Dataset statistics

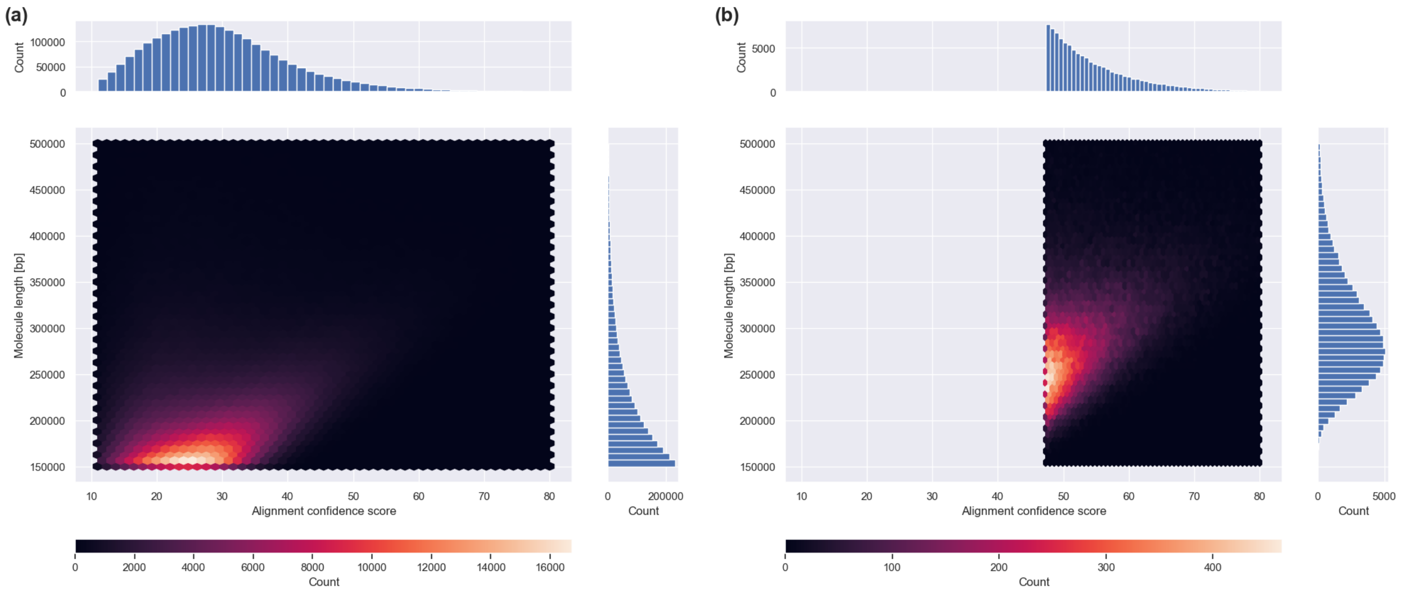


Figure S2: **Distribution of molecule length and confidence score**. The 2-d joint histogram of molecule lengths and confidence scores, along with their marginal histograms in the ground-truth set is shown. It can be seen that as expected, confidence grows with molecule length. On the left subplot (a) it is shown for the whole Bionano alignment output XMAP file, and on the right subplot (b), for the ground-truth set used in this work, chosen based on the 100K molecules with highest confidence score. From this ground-truth set the train, validation and tests subsets were sampled in this work, as described in subsection 2.3.

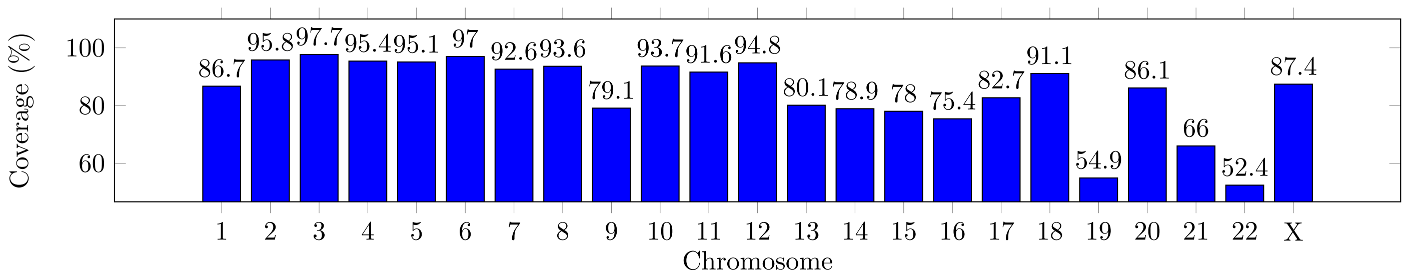


Figure S3: **Genome coverage of the molecules in the ground-truth set**. The human genome coverage of the ground-truth set of DNA molecules sampled for the analysis in this work (described in subsection 2.3) are shown. It can be seen that most of the chromosomes have a coverage of over 75%. The coverage was computed as follows: The union of the genome intervals for all molecules in the ground-truth set was computed after their alignment to the gnome (based on the Bionano's alignment output XMAP file, validated with DeepOM as described in subsection 2.3). This was computed per chromosome, and the ratio of the size of this union to the total size of the relevant chromosome reference sequence was calculated as the coverage. Such high coverage means that the test set we sampled from this ground-truth set will highly overlap with the training set genome-wise, but this is OK since we are interested in the model's ability to generalize to unseen images, since the genome is expected to be mostly the same in the training phase and then in the deployment of the model in actual use. So in practice we could train on data that has 100% genome coverage.

### S3 Alignment confidence and accuracy

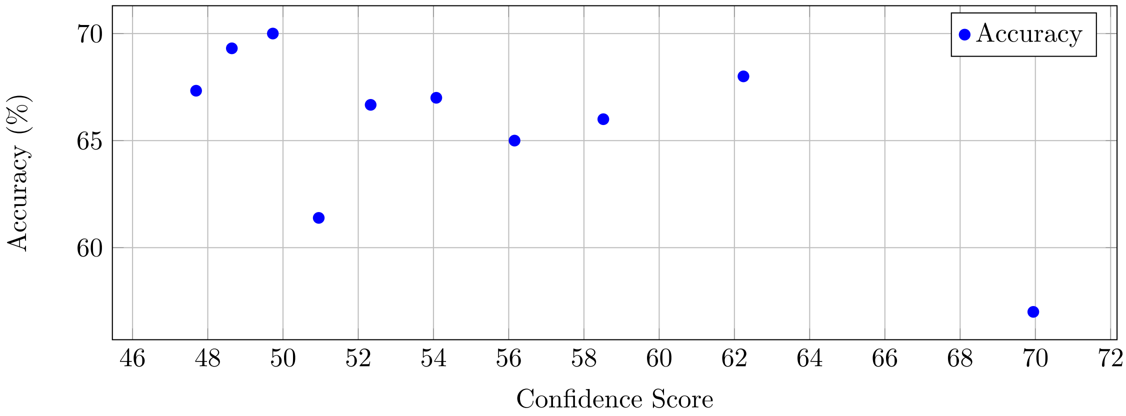


Figure S4: **Accuracy vs alignment confidence of ground-truth molecule**. In this figure, the accuracy of the model on cropped fragments of 50kb length is plotted against the median confidence of the subset of test-set molecules from which they were cropped. This was done to verify that the model will generalize well even to data of lower quality that could potentially have lower Bionano alignment confidence scores. The accuracy was computed as follows: The test set (that had 1000 ground-truth molecules, as described in subsection 2.3) was divided into 10 quantiles based on the confidence score of the molecules, each having 100 ground-truth long molecules. For each quantile subset of the test set, the model was evaluated on the cropped fragments of 50kb length from the long molecules in that quantile. The accuracy was computed as the ratio of the number of correctly mapped fragments to the total number of fragments in that quantile. The median confidence of the quantile subset was used as the x-axis value. As was done in the accuracy evaluation in the Results section 3

#### S4 Effect of retrieval K candidates count

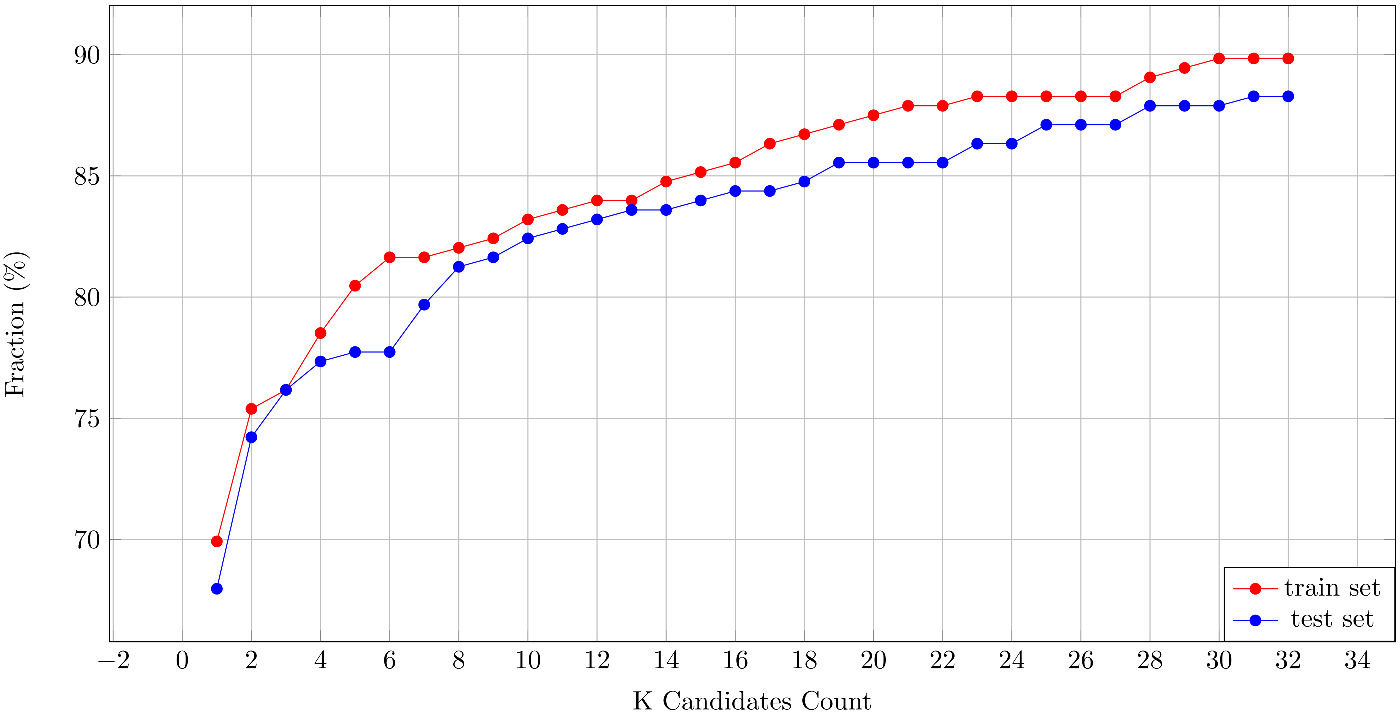


Figure S5: **The retrieval candidates K count**. The fraction of times the correct mapping is within the nearest K candidates is shown for 50 kb DNA fragments, for both the train and test subsets. The results were computed as follows. Given a subset of ground-truth molecules (train or test), 256 crops of 50 kb size were generated, in a similar way to Figure 6. For each crop, OM2Seq was used to retrieve the nearest 32 candidates from the genome reference, then for each K, the fraction of the 256 crops which had the correct mapping within the nearest K candidates (out of the 32) was computed. As expected, the performance on the train set is better, and also, increasing K increases the probability that the correct mapping is within the nearest K candidates. This reasoning motivated our integration of the combined approach of DeepOM + OM2Seq, as described in subsection 2.6, to improve the accuracy while still gaining the computational speedup of OM2Seq.
